# Deubiquitylase UCHL3 drives error correction at kinetochores and chromosome segregation independent of spindle assembly checkpoint

**DOI:** 10.1101/2020.03.31.018077

**Authors:** Katerina Jerabkova, Yongrong Liao, Charlotte Kleiss, Sadek Fournane, Matej Durik, Arantxa Agote-Arán, Laurent Brino, Radislav Sedlacek, Izabela Sumara

## Abstract

Equal segregation of chromosomes during mitosis ensures euploidy of daughter cells. Defects in this process may result in imbalance in chromosomal composition and cellular transformation. Two surveillance pathways, the spindle assembly checkpoint (SAC) and the error-correction (EC), exist at kinetochores that monitor microtubule attachment and faithful segregation of chromosomes at the metaphase to anaphase transition. However, the molecular understanding of the interplay between EC and SAC signaling remains limited. Here we describe a role of deubiquitylase UCHL3 in the regulation of EC pathway during mitosis. Downregulation or inhibition of UCHL3 leads to improper attachments of chromosomes to spindle microtubules and to chromosome alignment defects during metaphase. Frequent segregation errors during anaphase and consequently aneuploidy is also observed upon inactivation of UCHL3. Surprisingly, UCHL3 is not involved in SAC signaling as both recruitment of SAC proteins to kinetochores and timely anaphase onset are not perturbed in UCHL3-deficient cells. Mechanistically, UCHL3 interacts with and deubiquitylates the mitotic kinase Aurora B known to drive both SAC and EC signaling. UCHL3 promotes interaction of Aurora B with MCAK, important EC factor but does not regulate Aurora B binding to other interacting partners or subcellular localization of Aurora B. Our results thus suggest that UCHL3-mediated deubiquitylation functionally separates EC from SAC signaling during mitosis and is critical for maintenance of euploidy in human cells.

## Introduction

Faithful chromosome segregation during mitosis ensures genome stability and defects in this process may lead to imbalance in chromosomal composition named aneuploidy or polyploidy, which have been causally linked to cellular transformation (1). The kinetochore is a large proteinaceous structure built onto centromere regions of sister chromatids of all chromosomes where spindle microtubules attach, thereby driving faithful sister chromatid separation at the transition from metaphase to anaphase. At the kinetochore two surveillance mechanisms exist that monitor correct microtubule-kinetochore (MT-KT) attachments and bi-orientation of chromosomes: the spindle assembly checkpoint (SAC) (also known as the mitotic checkpoint) and the error-correction (EC) signaling (2). The SAC inhibits onset of chromosome segregation during anaphase until all chromosomes form proper attachments with the mitotic spindle (bi-orientation or amphitelic attachments), where each kinetochore is bound to microtubules emanating from the opposite spindle poles (3, 4). Unattached kinetochores create a binding platform for various SAC proteins including mitotic arrest deficient 2 (MAD2), budding uninhibited by benzimidazoles 3 (Bub3) and Bub1-related 1 kinase (BubR1) proteins, which form the Mitotic Checkpoint Complex (MCC) able to sequester the Cdc20 protein. Cdc20 represents the co-activator and the substrate specific adaptor of the E3-ubiquitin ligase Anaphase Promoting Complex/Cyclosome (APC/C), which ubiquitylates and targets Securin and Cyclin B proteins for 26S proteasomal degradation, thereby triggering anaphase onset (5). The kinetochore attachment defects such as syntelic (attached to single pole) or monotelic (mono-oriented) attachments, prevent SAC satisfaction and inhibit chromosome segregation, often leading to prolonged mitotic arrest and ultimately to cell death. Importantly, such defects in microtubule attachments to kinetochores can be repaired by the error-correction (EC) mechanism, which allows for selective destabilization of these erroneous configurations and for stabilization of bi-oriented attachments (6). One of the main factors driving error-correction is the mitotic kinase Aurora B (7, 8). Aurora B is a catalytic subunit of the so called Chromosomal Passenger Complex (CPC), which dynamically localizes to centromeres, kinetochores and microtubules during mitotic progression (9). Downregulation or inhibition of Aurora B stabilizes incorrect attachments (7, 8, 10–12) and many microtubule-associated Aurora B substrates were identified (13). For instance, activity of Aurora B is required for the kinetochore localization and function of mitotic centromere-associated kinesin (MCAK) (also known as Kif2C), which acts as a microtubule depolymerase on incorrectly formed MT-KT attachments (14–20). The recruitment of the kinesin CENP-E to kinetochores is regulated by Aurora B, playing an important role in the conversion from initial lateral MT-KT attachments to the end-on attachments and thereby in the bi-orientation of the chromosomes (11, 21). Conversion of the lateral to the end-on attachments as well as stabilization of bi-oriented attachments is also controlled by the Astrin-SKAP complex, which is antagonized by the action of Aurora B (22, 23). It has been initially suggested that the role of Aurora B in the regulation of SAC is indirect, namely that in the absence of spindle-exerted tension across the centromeres, the error-correction mechanism is activated leading to destabilization of erroneous MT-KT attachments and in turn to SAC potentiation (24, 25). However, accumulating evidence suggests the important direct role of Aurora B in SAC signaling (26) also through kinetochore recruitment of numerous SAC proteins (10, 11, 27–29). Thus, Aurora B is an upstream factor playing a critical role in both surveillance pathways SAC and EC during mitosis. However, the regulatory mechanisms able to functionally uncouple EC from SAC remain poorly characterized. Here, using unbiased screening approaches, we identify an unexpected role of the deubiquitylase UCHL3 which exerts specific function on EC independent of SAC signaling during mitosis of human cells.

## Materials and Methods

### Reagents and antibodies

TCID (4,5,6,7-tetrachlorodindan-1,3-dione) UCHL3 inhibitor (Ref. 27720-1), C9H2Cl4O2 was purchased from Tebu-Bio and used at 2μM working concentration. Monastrol (Ref. M8515) and 4′,6-Diamidino-2-phenylindole dihydrochloride (DAPI) (Ref. D8417) were purchased from Sigma-Aldrich. S-Trityl-L-cysteine (STLC), (Ref. ALX-105-011-M500) was purchased from Enzo Life Sciences and the proteasomal inhibitor MG132 (No. 1748) was purchased from Tocris bioscience. Protease inhibitors (cOmplete EDTA-free Protease Inhibitor Cocktail) were purchased from Roche.

UCHL3 antibody was produced by IGBMC antibody facility using immunized rabbits and purified with SulfoLink resins according to the manufacturer’s protocol. The Astrin polyclonal rabbit antibody was a kind gift from Ulrike Gruneberg (Cancer Research, UK). Commercial antibodies: Mouse monoclonal BubR1 (BD Biosciences, 612502 clone 9/BubR1), mouse monoclonal α-tubulin (Sigma Aldrich, T5168), human polyclonal CREST (Antibodies Incorporated, 15 234), rabbit polyclonal Aurora B (Abcam ab2254), mouse monoclonal UCHL3 (Sigma Aldrich, clone H7171), mouse monoclonal CENP-E (Thermo Scientific, MA1-5758), rabbit polyclonal GFP (Abcam, ab290)), Cyclin B1 (GNS1) (Santa-Cruz, sc-245), β-actin (Sigma, A2228), PP1γ (Santa Cruz, sc-517354), HOIP (Abcam, ab125189). MAD2L1 (Genetex, GTX104680, survivin (Abcam, ab469).

### Plasmids

All GFP plasmids used were cloned into pEGFP-N1 plasmid (Clontech): pEGFP-N1, pEGFP-N1-UCHL3-WT, pEGFP-N1-UCHL3-C/S. UCHL3 (NCBI, variant2 NM_006002.4) was cloned from human cDNA. UCHL3 catalytic dead mutant has cysteine 95 residue mutated to serine by G > C base exchange in the cysteine codon. For Immunoprecipitation experiments, N-terminally tagged pEGFP-Aurora B vector was used.

### Cell culture and synchronization of cells

All cell lines were purchased from ATCC and cultured at 37 °C in 5% CO2 humidified incubator, if not stated otherwise. HeLa Kyoto (human cervix carcinoma) cells were cultured in high glucose DMEM-GlutaMAX (4.5 g/L glucose) supplemented with 10% foetal calf serum (#9150), 1% penicillin and streptomycin. HeLa cells stably expressing Tubulin-GFP-H2B-mCherry were purchased from Ellenberg laboratory and a standard medium for HeLa cells was used to culture them. Human primary lung fibroblasts (IMR90) were cultured in EMEM containing non-essential AA, 2 mM L-glutamine, 1 mM sodium pyruvate, 1500 mg/L sodium bicarbonate, 10% foetal calf serum and gentamycin. Dld1-mCherry cell line was a kind gift from Don Cleveland and it was cultured in RPMI-1640 medium containing 2 mM L-glutamine, 10 mM HEPES, 1 mM sodium pyruvate, 4500 mg/L glucose, 1500 mg/L sodium bicarbonate, 10% fetal calf serum and gentamycin. Cells were trypsynized, counted in Neubauer chamber and seeded on 9-15 mm glass coverslips (Menzel-Glaser) in 24-well plates at a density 15 000 cells per well for all immunofluorescence experiments. To synchronize cells in prometaphase STLC (50 mM/DMSO) was diluted in medium to 5 μM and cells were treated 16 hours. Monastrol (100 mM/DMSO) was diluted to 100 μM and used for 16 hours. Monastrol washout: upon incubation cells were washed 5x with warm medium and released in fresh culture medium for different time periods as indicated in respective figures. Monastrol release: upon incubation cells were washed 5x with warm medium and released for 90 minutes to MG132 (50 μM) containing medium to arrest the cells in metaphase. Thymidine (200 mM/H2O) was used at 2 mM, cells were treated 16 hours, washed 3 times, released for 8 hours and treated again 16 hours with thymidine followed by 3 washes. Mitotic cells were observed 8.5 hours after the second release.

### Generation of stable cell lines

Stable cell lines were generated in HeLa Kyoto cells by random integration of GFP-UCHL3 plasmids. Three lines were generated: GFP-HeLa expressing empty GFP plasmid as a control, GFP-WT-UCHL3 expressing wild-type sequence of UCHL3 and GFP-C/S-UCHL3 expressing catalytic dead mutant of UCHL3. Expression levels of different proteins were estimated by western blot. Cell lines expressing near endogenous levels of UCHL3 proteins were chosen for immunoprecipitation experiments.

For retroviral-mediated silencing of genes induced by stable expression of short hairpin RNA (shRNA) targeting UCHL3 ATAGAAGTTTGCAAGAAGTTTA and a control sequence TAATCAGAGACTTCAGGCGG (targeting Firefly luciferase) was cloned into an LMP backbone. This plasmid contains long terminal repeats and retroviral packaging signal necessary for the virus production and PKG promoter driven expression of a cassette coding for puromycin resistance and GFP. Furthermore, for improved production of the shRNA, it contains a cassette with U6 promoter driven expression of miR30 microRNA context into which the designed shRNA sequences are cloned. Retrovirus was produced by transiently transfecting the Phoenix packaging cell line (G. Nolan, Stanford University, Stanford, CA) with the prepared plasmids. Supernatant was collected, filtered to remove cellular debris, polybrene was added and the supernatant was used to infect HeLa cells or Dld1 cells overnight. After 2 days, cells were selected using puromycin (1 mg/ml) for 48 hours. After selection, the presence of replicatively competent retrovirus was excluded using qPCR. Knockdown of UCHL3 was validated using qPCR.

### Generation of UCHL3−/− cell lines by CRISPR/Cas9 genome editing system

For generating HeLa UCHL3−/− cell lines, guide RNA (gRNA) was designed by online software Benchling (https://www.benchling.com/), 5′-CAAACAATCAGCAATGCCTG-3′ and 5′-TGAAGTATTCAGAACAGAAG-3′ and cloned into pX330-P2A-EGFP (Addgene) through ligation using T4 ligase (New England Biolabs). HeLa-Kyoto cells were transfected using Lipofectamine^™^ 2000 Transfection Reagent (Thermo Fisher Sceintific), 24 h after transfection, GFP-positive cells were collected by FACS (BD FACS Aria II) and seeded into 96-well plates. Obtained UCHL3−/− single-cell clones were examined by western blotting and sequencing of PCR-amplified targeted fragment by Phusion polymerase (Thermo Scientific). The PCR amplification primers were: 5′-CTGTAACGTGATCGTACAAA-3’ (forward) and 5′-GAATTAGAGCACCACCTACT-3′ (reverse).

### siRNA experiments

Cells were transfected by Oligofectamine at the final concentration 30 nM of the siRNA: UCHL3 siRNA-1 CAG CAU AGC UUG UCA AUA A, UCHL3 siRNA-2 GCA AUU CGU UGA UGU AUA U, UCHL3 3’UTR siRNA CUG CCA UAC ACU AAC UCA A, Ku80 siRNA (XRCC5 Dharmacon On-Target Plus) based on the manufacturer’s instructions. For rescue experiments cDNA and siRNA were co-transfected by Lipofectamine 2000 according to manufacturer’s instructions.

### Immunoprecipitations

HeLa cells stably expressing GFP-UCHL3 proteins were cultured, synchronized with STLC and harvested in 1 ml of lysis buffer (10 mM Tris HCl pH 7.5, 150 mM NaCl, 0.5 mM EDTA, 0.5% NP-40, protease inhibitor cocktail) per four 10 cm dishes in each condition. GFP-trap agarose beads (Chromotek) were blocked overnight in 3% BSA diluted in wash buffer (10 mM Tris HCl pH 7.5, 150 mM NaCl, 0.5 mM EDTA, protease inhibitor cocktail), washed three times in lysis buffer and incubated with 10 mg of cell extracts overnight, rotating at 4 degrees. Before elution, the beads were washed 5 times for 5 minutes with 1 ml washing buffer (centrifugation 500g, 2 minutes). The proteins were eluted from beads by boiling them in 2x Laemmli SDS sample buffer (BioRad) for 15 minutes. Samples were analyzed by western blotting.

For immunoprecipitation under denaturing conditions, UCHL3+/+ and UCHL3−/− HeLa cells were transfected with GFP-Aurora B or pEGFP-N1 and synchronized by monastrol for 18 h. Cells were lysed with Urea lysis buffer (8M Urea, 300 mM NaCl, 50 mM Na2HPO4, 50 mM Tris-HCl, 1 mM PMSF, pH 8) and sonicated, supernatants were cleared by centrifugation at 16000g for 15 minutes and incubated with GFP-Trap agarose beads (Chromotek) overnight at 4 °C. Beads were washed by Urea lysis buffer, eluted in 2 × Laemmli buffer and analyzed by western blotting.

### Western blot analysis

To isolate proteins, cells were scraped, pelleted by centrifugation at 4 °C and washed twice in PBS. Lysates were prepared using RIPA buffer (50 mM Tris-HCl pH7.5, 150 mM NaCl, 1% Triton X-100, 1 mM EDTA) supplemented with 1 mM NaF, 1 mM DTT and protease inhibitor cocktail Complete. Cells were lysed on ice by mechanical disruption with a needle (26G) and centrifuged at 10 000g for 30 minutes at 4 °C. Protein concentration of supernatant was measured by Bradford assay (Biorad). Samples were boiled for 10 minutes in Laemmli buffer with β-Mercaptoethanol (BioRad), resolved on 10% polyacrylamide gels or pre-cast gradient gels (Thermo Scientific, NW04120BOX) and transferred onto PVDF membrane (Millipore) using semi-dry transfer unit (Amersham). 5% non-fat milk was used for blocking and for antibody dilution. TBS-T (25 mM Tris-HCl, pH 7.5, 150 mM NaCl, 0.05% Tween) was used for washing.

### Quantitative PCR analysis

RNA was isolated from HeLa cells using kit from Machery Nagel and reversed transcribed by qScript® cDNA SuperMix (Quanta Bio) according to the protocols. For the qPCR 20 ng of cDNA were used per reaction and amplified with SybrGreen I master mix from Roche (04 887 352 001) at 60 °C and values were normalized to GAPDH. Human Ku80 primers were: Fwd GCTAATCCTCAAGTCGGCGT, Rev CAGCATTCAACTGTGCCTCG, GAPDH primers: Fwd CACCCAGAAGACTGTGGATGG, Rev GTCTACATGGCAACTGTGAGG.

### Immunofluorescence microscopy

Cells were washed once in PBS and fixed in 4% PFA for 17 minutes at room temperature (RT), washed three times in PBS and permeabilized in 0.5% NP-40 for 5 minutes, followed by three washes in PBS-T and blocking in 3% BSA in PBS-T for 90 minutes at RT or at 4 °C overnight. Primary antibodies were diluted in the blocking buffer (1:250) and incubated 2 hours at RT, washed 3 times 5 minutes in PBS-T and incubated with secondary antibodies at 1:5000 dilution for 1 hour at RT in the dark. Cells were washed 3 times 10 minutes in PBS-T and incubated with DAPI diluted in PBS at final concentration 1 μg/ml for 10 minutes at RT, followed by 2 washes in PBS-T. Cover slips were mounted on glass slides using Mowiol and dried overnight. High resolution images were taken using Leica spinning disc confocal microscope with Live SR module at 100x magnification and were processed in Image J followed by the analysis in Cell Profiler as described below.

For analysis of the microtubule motor proteins, prior to fixation, cells were incubated in extraction buffer (PHEM, 1 mM PMSF, 1 mM ATP, 0.5% Triton X-100) for 3 minutes at 37 °C and fixed after in 4% PFA for 2 minutes at 37 °C followed by two times 5 minutes incubation with 0,5% Triton in PBS at 37 °C. Cells were blocked in 3% BSA in PBS-T, followed the standard IF protocol from this step onwards. PHEM buffer (pH 6.9, PIPES 45 mM, HEPES 45 mM, EGTA 10 mM, MgCl2 5 mM).

### Live video imaging

Cells were seeded in a glass bottom dish, 30 mm diameter with 4 compartments and synchronized with monastrol or DTBR. For acquisition, Leica CSU-W1 spinning disk, 63x objective, water immersion was used with time frame 5 min and z step 2μm. Cells were placed into a humid heated chamber with 5 % CO2 and 80 % humidity in the microscope and in case of monastrol release, cells were rinsed 5 times with warm medium to wash out the monastrol. Eight positions were selected for each condition and pictures were taken during 4 hours. Images were processed in ImageJ and maximum projections were created.

To quantify percentage of cells with misaligned chromosomes, cells were stained with DAPI and the experiment was performed in a blinded setup. For each category, the number of cells having aligned and misaligned chromosomes was counted and the percentage of total cells was quantified. In each experiment between 200-800 cells per condition were counted.

### Cell Profiler data analysis

Cell Profiler 3.1.8 was used to analyze the immunofluorescent images. Irregular nuclei: Form factor was calculated as 4*π*Area/Perimeter2. Primary objects (nuclei) were identified based on the DAPI channel and the form factor for each nucleus was measured in all conditions. Threshold was set to discriminate between regular and irregular nuclei and the percentage of irregular nuclei was calculated. Relative kinetochore intensity was quantified based on immunofluorescence image of CREST staining (kinetochore regions). The mean intensity of CREST and protein of interest (POI): BubR1, Astrin, Aurora B, MCAK was measured in the selected area and the ratio of the POI intensity to the CREST intensity (POI/ CREST) was calculated and the values were normalized to the control (control = 1). Spatial attributes were assigned manually to the individual chromosomes (C- centered, P- polar). Two focus planes were chosen from each Z-stack.

### Statistics

To statistically evaluate the data, parametric One-Sample T-test was used for samples normalized to the control and parametric Two-sample T-test was used for the rest of the conditions. Values were considered significantly different if p < 0.05 and stars were assigned as follows: p <0.05 *, p <0.01 **, p <0.001 *** p < 0.0001 ****. In all graphs, results are shown as mean ± SEM (or SD) of at least three independent experiments. Details for each graph are listed in figure legends.

## Results

### UCHL3 controls euploidy of human cells

To identify factors that control euploidy of human cells, we performed a high content visual siRNA screen in HeLa cells for approximately 100 known and predicted human deubiquitylases (DUBs) using established approach previously described for the ubiquitin-binding proteins (30). We scored for terminal phenotypes of multilobed nuclei and multinucleation often resulting from defects in chromosome segregation and cytokinesis such as those observed upon downregulation of Aurora B and UBASH3B (30) (Fig. 1A). Downregulation of the top DUB hit (Table S1), Ubiquitin carboxyl-terminal hydrolase isozyme L3 (UCHL3) showed similar strong nuclear atypia in the screen relative to the positive controls (Aurora B and UBASH3B) (Fig. 1A). UCHL3 belongs to the family of ubiquitin C-terminal hydrolases (UCH) containing also UCHL1, BRCA1 associated protein 1 (BAP1) and UCHL5. UCHL3 has been implicated in the neurodegenerative disorders (31) and cancer (32, 33) but it has not been linked to the regulation of mitosis. To confirm its potent role in cell division, we downregulated UCHL3 by specific siRNAs used in the secondary screen (Table S1) (Fig. 1B-D), and a distinct siRNA targeting a 3’UTR region of UCHL3 gene (Fig. 1E-G). In agreement with results obtained by the unbiased screening (Fig. 1A), downregulation of UCHL3, but not UCHL1, led to strong reduction in its protein levels (Fig. 1B-G) and markedly increased the number of cells with irregular, multilobed nuclei (Fig. 1B, D, E, F). Treatment with potent, cell permeable UCHL3-specific inhibitor 4,5,6,7-Tetrachloro-1*H*-Indene-1,3(2*H*)-dione (TCID) likewise resulted in accumulation of cells with nuclear morphology defects (Fig. 1H, I), suggesting that UCHL3 DUB activity might regulate faithful cell division and control euploidy of human cells.

**Fig. 1.**
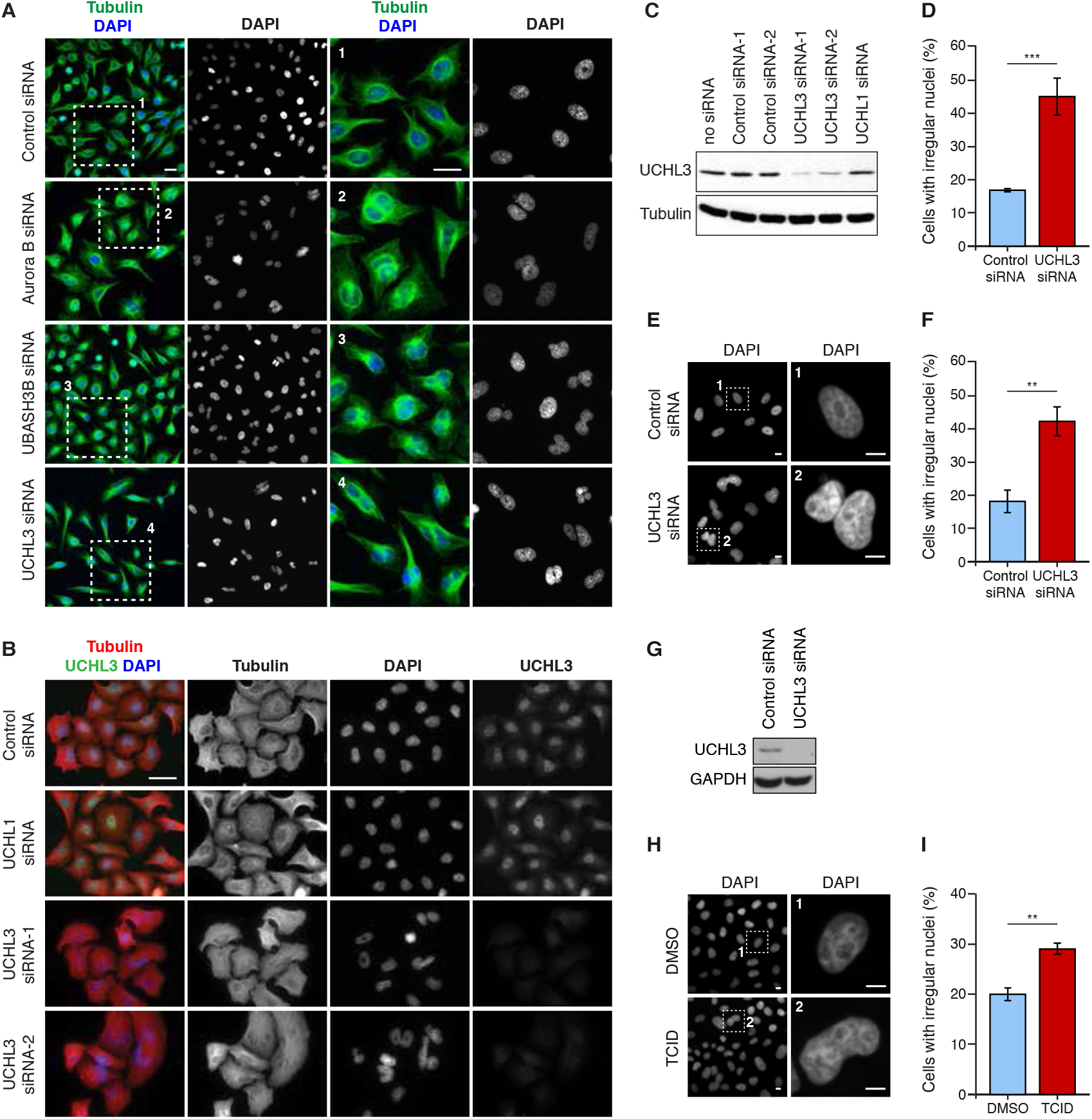
UCHL3 controls euploidy of human cells. UCHL3 was downregulated by specific siRNAs used in the secondary screen (Table S1) (A-D), and a distinct siRNA targeting a 3’UTR region of UCHL3 gene (E-G). (A) HeLa cells were treated with indicated siRNA pools and analyzed by immunofluorescence microscopy. The magnified regions are shown in the corresponding numbered panels, scale bar 15μm. (B-D) HeLa cells were transfected with single siRNAs as indicated and analyzed by immunofluorescence (B), scale bar 20μm and by (C) western blotting. (D) The percentage of cells with irregular nuclei was quantified, mean ± SD of 3 experiments. Control = 16.4 ± 0.2, UCHL3 siRNA-1 = 44.5 ± 5.3, p=0.0008. (E-G) HeLa cells were treated with control and UCHL3 3’UTR siRNA as indicated and analyzed by immunofluorescence (E), scale bar 10μm. The magnified regions are shown in the corresponding numbered panels, scale bar 5μm. (F) Percentage of cells with irregular nuclei, mean of 4 experiments ± SEM. Control = 18.3 ± 3.3, n= 886 cells, UCHL3 3’UTR siRNA = 42.3 ± 4.3, n = 828 cells, p = 0.0043. (G) Western blot analysis confirming UCHL3 knockdown. (H-I) HeLa cells were with treated UCHL3 inhibitor (TCID) and analyzed by immunofluorescence (H), scale bar 10μm. The magnified regions are shown in the corresponding numbered panels, scale bar 10μm. (I) Percentage of cells with irregular nuclei, mean of 4 experiments ± SEM. Control (DMSO) = 19.5 ± 1.1, n = 2110 cells, TCID = 28.6 ± 1.1, n = 2230 cells, p = 0.0012.

### UCHL3 regulates fidelity but not timing of chromosome segregation

To understand if and how UCHL3 regulates mitotic progression, we employed spinning disk live video microscopy of HeLa cells stably expressing histone marker H2B-mCherry which were synchronized in prometaphase using an Eg5 inhibitor monastrol and released. This analysis showed that downregulation of UCHL3 led to chromosome alignment defects in metaphase (Fig. 2A) and to segregation errors in anaphase (Fig. 2A-C). Interestingly, the time of mitotic progression from prometaphase to anaphase was largely unaffected by downregulation of UCHL3 (Fig. 2D). The same results were obtained in the p53 negative DLD1 colorectal adenocarcinoma cell line stably expressing histone marker H2B-mCherry where downregulation of UCHL3 (Fig. S1E) led to strong segregation defects but not to statistically significant delay in the anaphase onset (Fig. 2E-H). To exclude any indirect effects of monastrol usage, HeLa H2B-mCherry and DLD1 H2B-mCherry cell lines were synchronized in G1/S by a double thymidine block and release protocol and mitotic progression was analyzed using spinning disk confocal live imaging. Also under these experimental conditions, downregulation of UCHL3 led to alignment and segregation defects in both cell lines without affecting the time from nuclear envelope breakdown to anaphase (Fig. S1A-D). Moreover, CRISPR-CAS9-mediated knock-out cells of UCHL3 (Fig. 5A), which were synchronized in G1/S by a double thymidine block and release protocol progressed normally through cell cycle relative to isogenic control cells as monitored by Cyclin B accumulation and degradation (Fig. S1F). Likewise, no differences in protein levels of HOIP the subunit of the linear chain assembly E3 ligase LUBAC (34) or Protein phosphatase 1 gamma (PP1*γ*) (35) both critically involved in alignment of chromosomes were observed in UCHL3 knock-out cells synchronously progressing through cell cycle relative to controls (Fig. S1F). These results suggest that UCHL3 regulates fidelity but not timing of chromosome segregation during mitosis. Chromosome segregation errors observed in UCHL3-deficient cells are likely to induce NoCut checkpoint and/or inhibit abcission and cytokinesis (36, 37) leading to irregular nuclei phenotype (Fig. 1A, B, E, H).

**Fig. 2.**
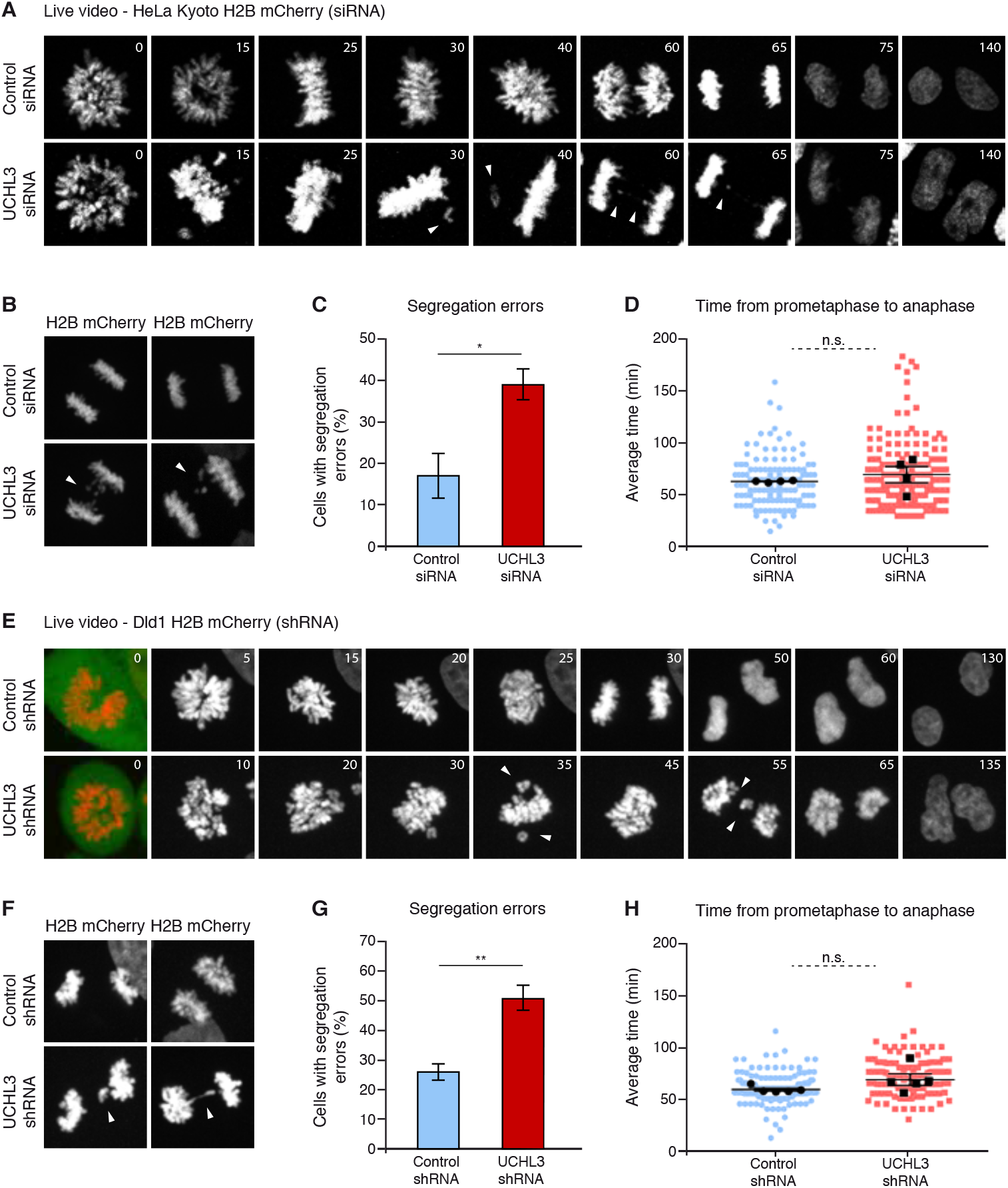
UCHL3 regulates fidelity but not timing of chromosome segregation. HeLa cells stably expressing H2B-mCherry (DNA) were treated with siRNA as indicated and synchronized in prometaphase, t=0 represents release from monastrol. (A) Selected time-frames showing merge of tubulin (green), DNA (red) signal and DNA single channel (gray). (B) Examples of observed segregation errors. (C) Percentage of cells with segregation errors, mean ± SEM of 4 experiments. Control = 17.07 ± 5.92, n= 127 cells, UCHL3 siRNA 39.01 ± 3.78, n= 198 cells, p= 0.0354. (D) Average time needed to proceed from prometaphase to anaphase, mean [min] ± SEM of 4 experiments. Control = 63.8 ± 0.4, UCHL3 siRNA = 70.4 ± 7.0, p= 0.4450. Colored dots represent values of individual cells, black symbols represent the mean experimental value. (E) Selected time frames of Dld1 cells stably expressing H2B-mCherry and GFP-shRNA as indicated. Cells were synchronized to prometaphase, t=0 represents release from monastrol. (F) Examples of observed segregation errors. (G) Percentage of cells with segregation errors, mean ± SEM of 5 experiments. Control = 26.1 ± 3.2, n= 125 cells, UCHL3 shRNA = 50.7 ± 5.4, n= 118 cells, p= 0.0069. (H) Average time needed to proceed from prometaphase to anaphase, mean [min] ± SEM of 5 experiments. Control = 59.2 ± 1.2, UCHL3 shRNA 68.7 ± 4.5, p= 0.1299. Colored dots represent values of individual cells, black symbols represent the mean experimental value.

### UCHL3 does not regulate SAC signaling

The lack of effects on the mitotic timing upon UCHL3 downregulation is surprising because errors in chromosome alignment should be detected by the SAC machinery and delay the onset of anaphase until all chromosomes are bi-oriented. To understand if UCHL3 contributes to SAC signaling, we performed immunofluorescence microscopy to analyze kinetochore localization of SAC protein BubR1. Downregulation of UCHL3 did not affect the localization of endogenous BubR1 to kinetochores in prometaphase (Fig. S2A-C). In addition, protein levels of BubR1 and MAD2 were not changed throughout the cell cycle in the UCHL3 knock-out cells relative to controls (Fig. S1F). We conclude that UCHL3 does not regulate recruitment of SAC protein BubR1 to kinetochores or its stability which could explain the results on the mitotic timing (Fig. 2D, H and Fig. S1C, D).

### UCHL3 activity specifically regulates EC pathway and chromosome bi-orientation

EC pathway represents the second surveillance mechanism ensuring chromosome bi-orientation and fidelity of chromosome segregation. To understand if and how UCHL3 contributes to EC signaling, we synchronized cells in metaphase by monastrol block and release into proteasome inhibitor MG132. Downregulation of UCHL3 led to dramatic increase of metaphase cells with misaligned chromosomes often detected in the proximity of the spindle poles (Fig. 3A, B). To understand if UCHL3 specifically regulates chromosome alignment to metaphase plate we performed rescue experiments, where 3’UTR directed siRNAs to UCHL3 gene were simultaneously transfected with GFP-tagged wild type or catalytically inactive versions of UCHL3 expressed at nearly endogenous levels (Fig. 3C, D). Expression of wild-type, but not catalytically inactive DUB UCHL3, rescued chromosome alignment defects upon downregulation of UCHL3 (Fig. 3C, E). These results suggest that UCHL3 specifically regulates chromosome bi-orientation in a manner that is dependent on its catalytic activity. These observations were confirmed in CRISPR-CAS9-mediated knock-out cells of UCHL3 (Fig. 5A), which also displayed chromosome alignment defects (Fig. 3F, 6E). Interestingly, downregulation of Ku80, reported direct deubiquitylation substrate of UCHL3 (38) and the core non-homologous end-joining (NHEJ) factor (Fig. S3A), did not result in defects in chromosome alignment as compared to UCHL3 siRNA and relative to control siRNA treated cells (Fig. S3B, C), suggesting that mitosis-specific regulation of chromosome alignment by UCHL3 cannot be attributed to regulation of Ku80. To further exclude the possibility that chromosome alignment errors observed in UCHL3-deficient cells are indirect due to possible defects in earlier cell cycle stages, we used UCHL3-specific inhibitor TCID, which led to the similar nuclear morphology defects (Fig. 1H, I). We compared longer treatments where the inhibitor was used simultaneously with monastrol for 16 hours and after the release into MG132 to allow for the formation of metaphase plates with the short treatment where TCID was only administered after monastrol release into MG132 (Fig. 3G). Importantly, the results from long (Fig. 3H, I) and short (Fig. 3J) TCID treatments were almost indistinguishable and both experimental conditions led to accumulation of misaligned polar chromosomes relative to solvent control. The same results were obtained in human primary lung fibroblasts IMR-90 (Fig. 3K-M). Our results thus demonstrate that UCHL3 activity is specifically required for the correction of improperly aligned chromosomes as cells progressively form the metaphase plate.

**Fig. 3.**
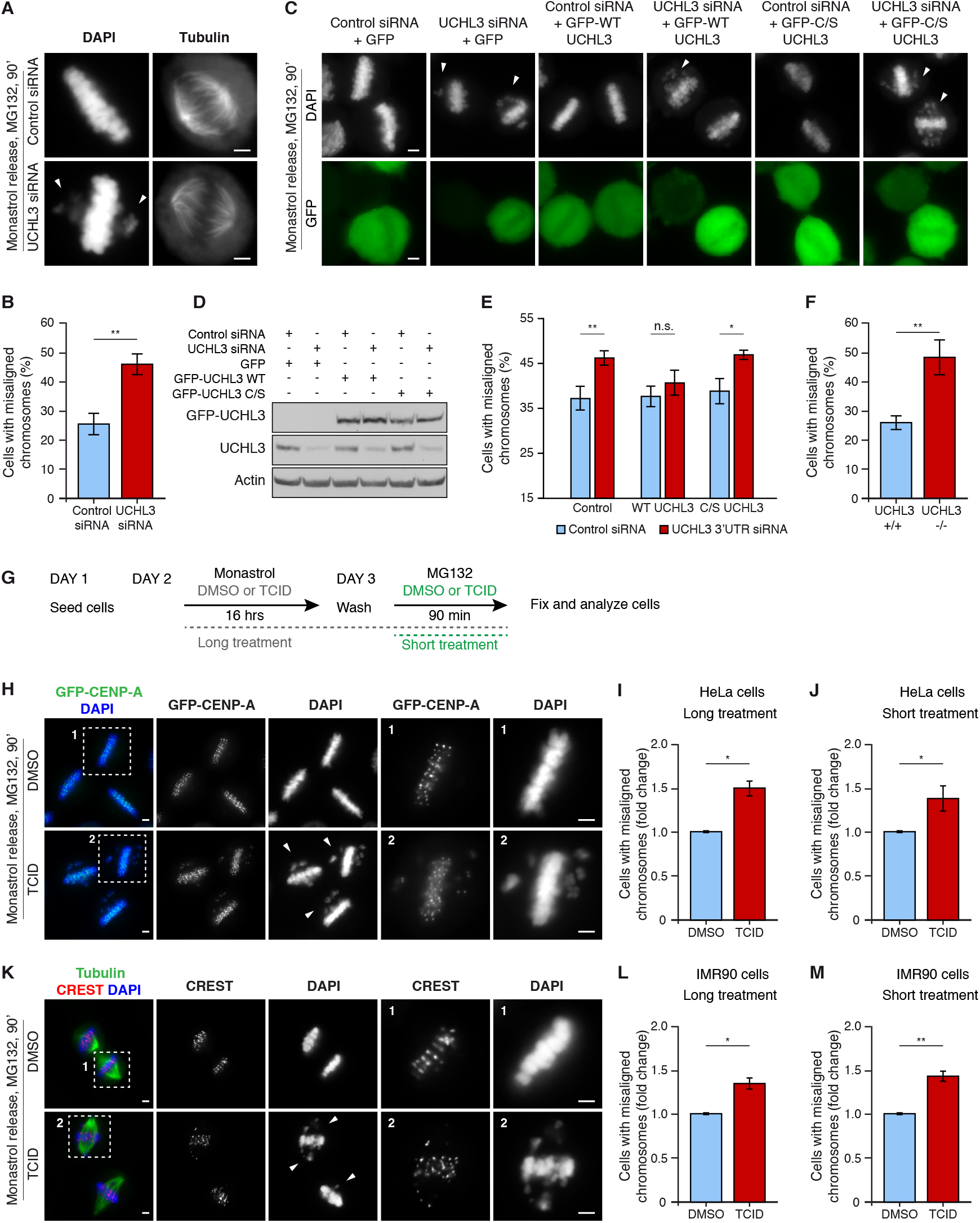
UCHL3 activity specifically regulates EC pathway and chromosome bi-orientation. (A) HeLa cells were treated with indicated siRNAs, synchronized to metaphase and analyzed by immunofluorescence, scale bar 2μm. (B) Percentage of cells having at least one misaligned chromosome, mean ± SEM of 7 experiments. Control = 25.2 ± 3.7, n = 2356 cells, UCHL3 3’UTR siRNA = 45.4 ± 4.2, n = 2801 cells, p = 0.0034. (C) HeLa cells co-transfected with siRNA as indicated and GFP, GFP-WT-UCHL3 (wild type), GFP-C/S-UCHL3 (catalytic dead) proteins were synchronized to metaphase and analyzed by immunofluorescence, scale bar 2μm. (D) UCHL3 knockdown efficiency and expression of the GFP-forms of UCHL3 were analyzed on the same membrane by western blotting with antibodies against UCHL3 and Actin. (E) Percentage of cells with misaligned chromosomes was quantified for each category, mean ± SEM of 4 experiments, 37.3 ± 2.5, 46.2 ± 1.7, 37.5 ± 2.6, 40.9 ± 3.0, 39.0 ± 2.7, 46.7 ± 0.9 with total number of cells analyzed: 1670, 2632, 1943, 2088, 2266, 2149 and p values p = 0.0242, p = 0.4306, p = 0.0366, respectively. (F) UCHL3 wild type (UCHL3+/+) and knockout (UCHL3−/−) HeLa cells were synchronized to metaphase and the percentage of misaligned chromosomes was quantified, mean ± SD of 3 experiments, UCHL3+/+ = 23.9 ± 2.5, UCHL3−/− = 47.6 ± 6.0, p = 0.0033. (G) Synchronization protocol for ‘long treatment’ (gray) and ‘short treatment’ (green). (H) HeLa cells stably expressing GFP-CENP-A were synchronized by long-term treatment protocol described in G and analyzed by immunofluorescence. Magnified regions are shown in the corresponding numbered panels, scale bar 2μm. (I) Cells with misaligned chromosomes were quantified, shown as fold change, mean ± SEM of 3 experiments. Control (DMSO) = 1, n= 830 cells, TCID = 1.50 ± 0.06, n = 1090 cells, p = 0.0157. (J) Quantification of misaligned chromosomes in short-term treatment shown as the fold increase, mean ± SEM of 4 experiments. Control (DMSO) = 1.0, n = 1260 cells, TCID = 1.39 ± 0.13, n = 1541 cells, p= 0.0394. (K) Human primary fibroblasts (IMR90) were analyzed as described in H. (L) Cells with misaligned chromosomes after a long-term treatment were quantified, shown as fold change, mean ± SEM of 3 experiments. Control (DMSO) = 1, n= 1378 cells, TCID = 1.34 ± 0.07, n = 1571 cells, p = 0.0190. (M) Quantification of misaligned chromosomes after a short-term treatment shown as the fold increase, mean ± SEM of 5 experiments. Control (DMSO) = 1.0, n = 951 cells, TCID = 1.43 ± 0.06, n=994 cells, p= 0.0055. Arrowheads point to misaligned chromosomes.

### UCHL3 is required for the formation of stable MT-KT attachments

To understand if UCHL3 is involved in the formation or maintenance of stable MT-KT attachments at the prometaphase to metaphase transition, we analyzed kinetochore localization of Astrin, the marker of stable amphitelic attachments in metaphase-synchronized cells. Downregulation of UCHL3 markedly reduced the recruitment of Astrin to kinetochores on centered chromosomes located in the proximity to the metaphase plate as well as on polar chromosomes located closer to the spindle poles (Fig. 4A-C) relative to control cells, suggesting that UCHL3 may be directly involved in the formation of proper MT-KT attachments. Based on these observations, we conclude that UCHL3 is involved in the EC machinery ensuring chromosome bi-orientation at mitotic spindle.

**Fig. 4.**
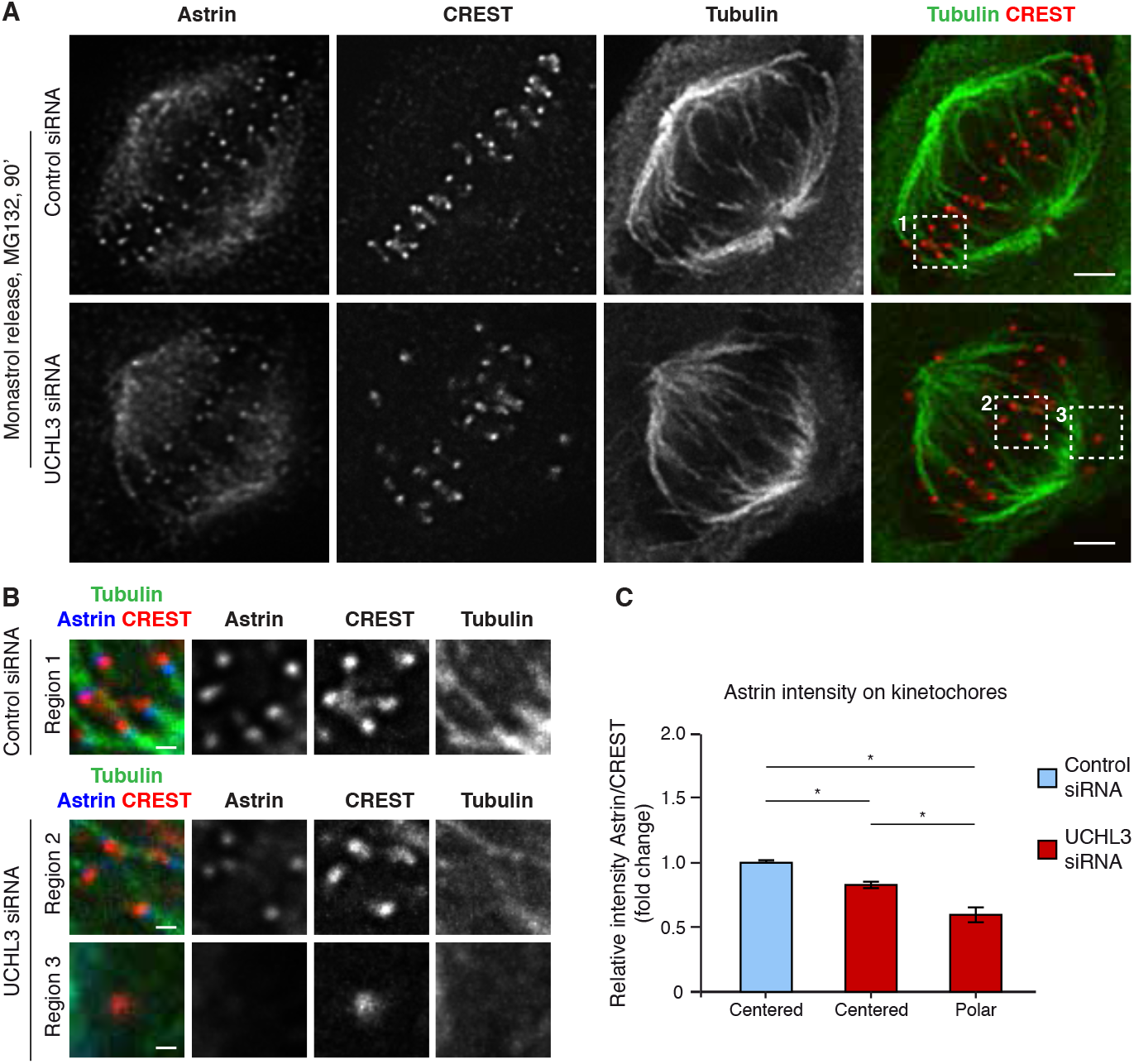
UCHL3 is required for the formation of stable MT-KT attachments. (A) HeLa cells were treated with siRNA as indicated, synchronized to metaphase and analyzed by immunofluorescence, scale bar 2 μm. (B) Magnified regions corresponding to numbered boxes in A, scale bar 1μm. (C) Relative Astrin intensity (Astrin/CREST ratio) on kinetochores of centered and polar chromosomes was quantified and is shown as a fold change to control. Results are presented as mean ± SD of 4 experiments. Control = 1, n= 4017 kinetochores, UCHL3 siRNA Centered (KD_C) = 0.82 ± 0.06, n = 3962 kinetochores, UCHL3 siRNA Polar (KD_P) = 0.59 ± 0.07, n = 470 kinetochores, p values: Control to KD_C p= 0.0114, control to KD_P p = 0.0107, KD_C to KD_P p = 0.0244.

### UCHL3 interacts with and deubiquitylates Aurora B in mitotic cells

Since UCHL3 regulates EC machinery during prometaphase, we hypothesized that UCHL3 acts on Aurora B, one of the major upstream factors driving EC pathway. Interestingly, Western Blotting analysis of UCHL3 knock-out cells revealed no differences in total protein levels of Aurora B (Fig. 5A, B) relative to control cells but demonstrated a smear of slower migrating forms of Aurora B detected in prometaphase and in early time points (15 minutes) after monastrol release specifically in UCHL3 knock-out cells (Fig. 5A). No differences in protein levels of Cyclin B, PP1*γ* and CENP-E were observed in the absence of UCHL3 relative to control cells during progression through prometaphase (Fig. 5B). No changes of protein levels of Aurora B were also observed in UCHL3 knock-out cells released from G1-arrest synchronously progressing through cell cycle relative to controls (Fig. S1F). Moreover, immunoprecipitated GFP wild-type could visibly interact with the endogenous slower migrating forms of Aurora B and with ubiquitin (Fig. 5C) in prometaphase cells, suggesting that UCHL3 may interact with polyubiquitin-modified forms of Aurora B. Unexpectedly, catalytically inactive UCHL3 appeared to interact with the unmodified Aurora B to a lesser extend (Fig. 5C), a result which requires further investigation. Immunoprecipitation of GFP-Aurora B under denaturing conditions confirmed existence of polyubiquitin-modified forms of Aurora B in prometaphase cells (Fig. 5D). Importantly, in the UCHL3 knock-out cells strong increase of polyubiquitin modification on immunoprecipitated GFP-Aurora B was observed (Fig. 5D). These results are consistent with the possibility that UCHL3 deubiquitylates Aurora B in prometaphase cells to specifically regulate its function in EC pathway.

**Fig. 5.**
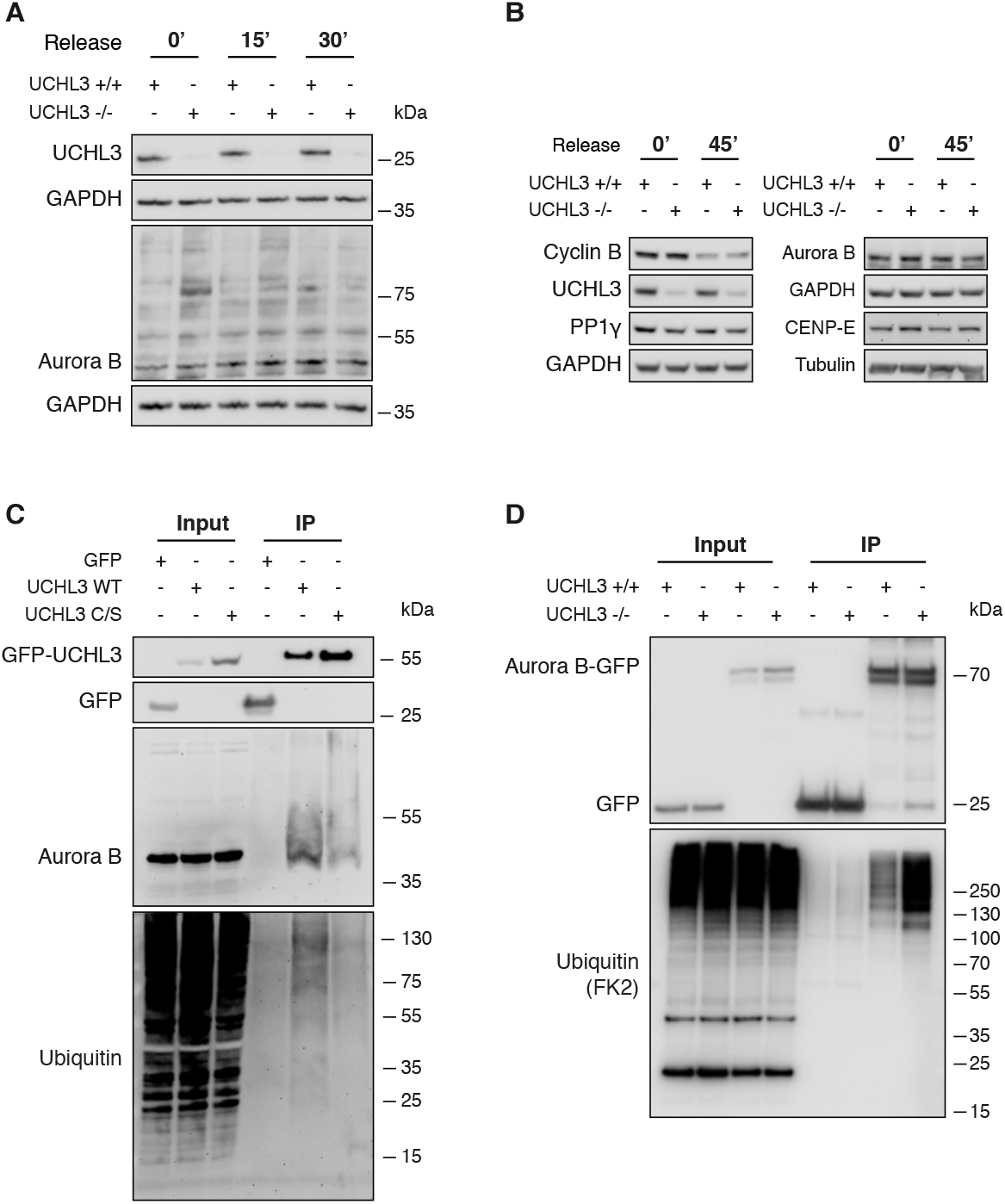
UCHL3 interacts with and deubiquitylates Aurora B in mitotic cells. (A, B) UCHL3+/+ and UCHL3−/− HeLa cells were synchronized in prometaphase by monastrol, released into fresh medium for 15, 30 or 45 minutes and analyzed by western blotting. (C) HeLa cells stably expressing, GFP, GFP-UCHL3-WT (wild type) or GFP-UCHL3-C/S (catalytic dead) proteins were synchronized in prometaphase (STLC) and the lysates were immunoprecipitated (IP) with GFP-Trap beads and analyzed by western blotting. (D) UCHL3+/+ and UCHL3−/− HeLa cells were transfected with Aurora B-GFP plasmid, synchronized in prometaphase (monastrol), collected for IP with GFP-Trap beads under denaturing conditions and analyzed by western blotting. Membranes were probed with GFP and FK2 antibody against conjugated ubiquitin.

### UCHL3 promotes interaction of Aurora B with EC factor MCAK

Since UCHL3 does not regulate protein levels or degradation of Aurora B (Fig. 5A, B and Fig. S1F), we next aimed at understanding possible molecular outcomes of UCHL3-mediated deubiquitylation of Aurora B. Ubiquitylation of Aurora B has been implicated in the regulation of its localization to mitotic structures (30, 39–42), we therefore first analyzed and quantified Aurora B chromosomal intensities in control and UCHL3-deficient cells. No differences of Aurora B signal could be observed relative to control cells (Fig. 6A, B). These observations were confirmed in UCHL3 knock-out cells (Fig. 6E), which displayed chromosome alignment defects (Fig. 3F). Likewise, analysis of intensity and kinetochore localization of MCAK, one of the main EC factors and direct substrates of Aurora B (15, 18), in UCHL3 deficient cells revealed no differences relative to control cells (Fig. 6A, C, D). Interestingly, we observed reduced interaction of immunoprecipitated GFP wild-type Aurora B with MCAK in UCHL3 knock-out cells relative to control cells while the binding of Aurora B to the CPC component Survivin and kinetochore factor Astrin remained unchanged in UCHL3 knock-out cells (Fig. 6F). These results suggest that UCHL3 activity promotes interaction of Aurora B with MCAK during chromosome alignment in prometaphase.

**Fig. 6.**
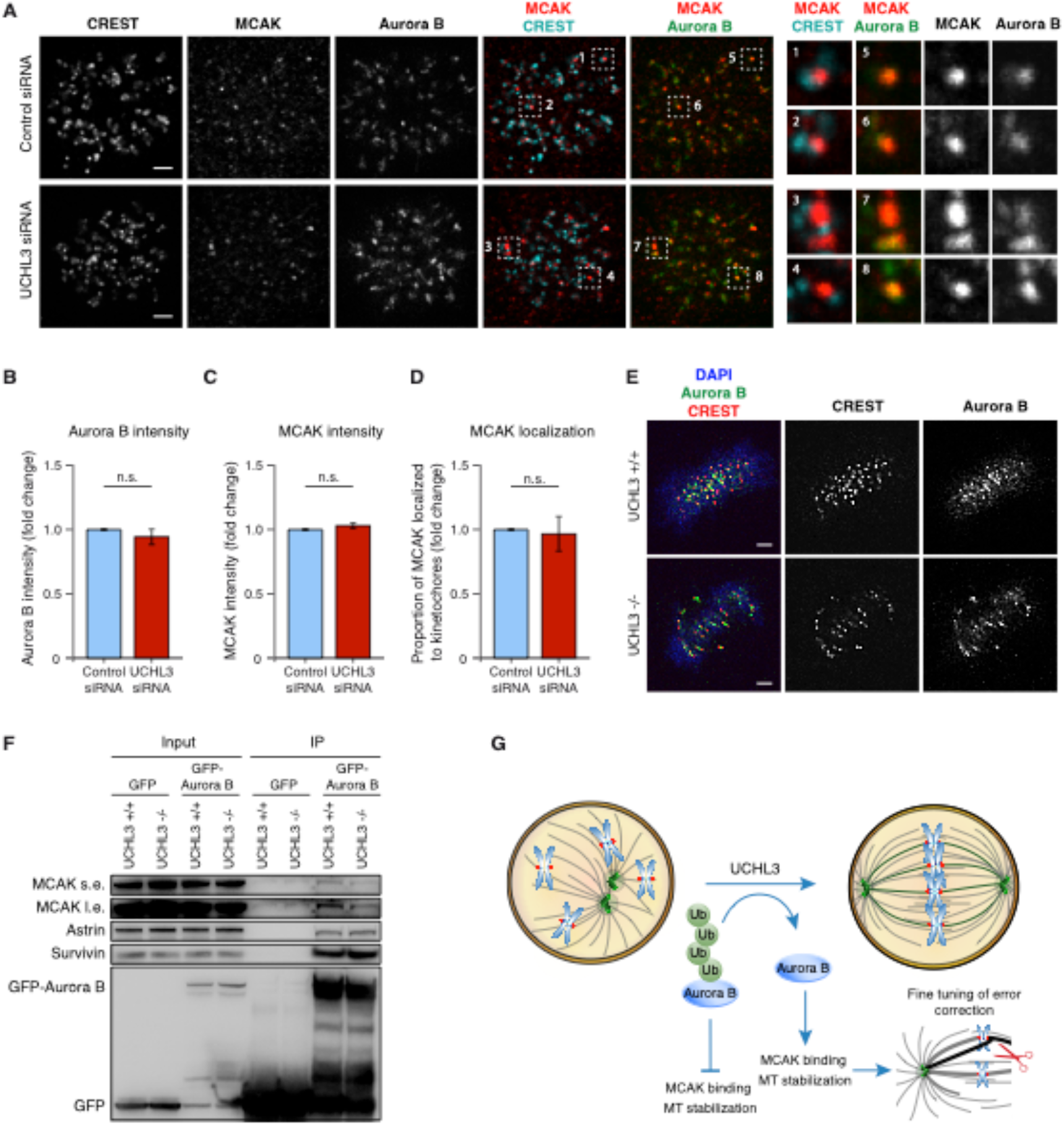
UCHL3 promotes interaction of Aurora B with MCAK. (A) HeLa cells stably expressing Aurora B-GFP were treated with siRNA as indicated, synchronized to prometaphase and analyzed by immunofluorescence with antibodies against MCAK and CREST, maximum intensity projections, scale bar 2μm. Right: Magnified regions corresponding to numbered boxes. (B) Total Aurora B intensity was quantified from single Z and normalized to control, mean of 5 experiments ± SEM. Control= 1, n= 88 cells, UCHL3 siRNA = 0.946 ± 0.054, n= 115 cells, p= 0.3775. (C) Total MCAK intensity from single Z was quantified and normalized to control, mean of 3 experiments ± SEM. Control= 1, n=73 cells, UCHL3 siRNA = 1.027 ± 0.024, n= 100 cells, p= 0.4967. (D) Percentage of MCAK signal on kinetochores normalized to control, mean of 3 experiments ± SEM. Control= 1, n= 4944 kinetochores, UCHL3 siRNA= 0.95 ± 0.05, n= 5460 kinetochores, p= 0.4992. (E) UCHL3+/+ and UCHL3−/− HeLa cells synchronized to metaphase and analyzed by immunofluorescence with antibodies against Aurora B and CREST, scale bar 2μm. (F) UCHL3+/+ and UCHL3−/− HeLa cells were transfected with Aurora B-GFP plasmid, synchronized by monastrol, collected for IP with GFP-Trap beads and analyzed by western blotting. The panel shows representative result of three independent experiments. (G) A hypothetical model how UCHL3 controls chromosome alignment and segregation. UCHL3 interacts with Aurora B in prometaphase and deubiquitylates it, which is necessary for proper Aurora B function in correction of erroneous MT-KT attachments. UCHL3-mediated deubiquitylation of Aurora B may promote its binding to MCAK and thus microtubule stabilization thereby fine-tuning EC machinery as cells progressively form metaphase plate. Polyubiquitylated Aurora B cannot interact with MCAK, resulting in microtubule destabilization, alignment and segregation errors. Green circle (Ub) = ubiquitin, blue oval = Aurora B kinase.

## Discussion

Taken together, our data suggest a model (Fig. 6G) how UCHL3-mediated deubiquitylation maintains euploidy in human cells. UCHL3-deficient cells display strong defects in bi-orientation of chromosomes at the mitotic spindle during metaphase (Fig. 3A-F) and frequent errors in chromosome segregation during anaphase (Fig. 2A-C, E-G and Fig. S1A, B). It is highly probable that nuclear morphology defects and aneuploidy observed in the absence of UCHL3 (Fig. 1A, B, D-F, H, I) result from an abscission block caused by lagging chromosomes during anaphase (36, 37). It would be interesting to know if the lagging chromosomes observed during anaphase of UCHL3-deficient cells correspond to the same chromosomes which failed to align at the metaphase plate. Indeed, our data demonstrate that in the absence of UCHL3 chromosomes with incorrect MT-KT attachments accumulate at the transition from prometaphase to metaphase where kinetochore localization of Astrin, a marker of stable MT-KT attachment (Fig. 4A-C) is perturbed. Our results suggest that UCHL3 specifically regulates alignment of chromosomes as cells progress from prometaphase to metaphase. First, the alignment defects observed upon UCHL3 downregulation by siRNA can be rescued by expression of WT but not catalytically inactive form of UCHL3 (Fig. 3 C-E). Similarly, CRISPR-CAS9-mediated UCHL3 knock-out cells display unaligned polar chromosomes in metaphase relative to controls (Fig. 3F). In addition to HeLa cells, the alignment defects can also be observed in UCHL3-deficient p53 negative DLD1 cells (Fig. 2E, Fig. S1B) and in human primary lung fibroblasts IMR-90 (Fig. 3K-M). It is unlikely, that observed alignment defects are due to misregulation of DNA repair by the NHEJ factor Ku80, a direct deubiquitylation substrate of UCHL3 (38), as Ku80-deficient cells display no differences in number of misaligned chromosomes relative to control cells in contrast to UCHL3-downregulated cells (Fig. S3A-C). Finally, the use of UCHL3-specific inhibitor TCID for a short period during prometaphase led to accumulation of misaligned polar chromosomes relative to solvent control, a defect almost indistinguishable from longer TCID treatments. Thus, UCHL3 activity is specifically required for the correction of improperly aligned chromosomes as cells progressively form the metaphase plate.

Several lines of observations suggest that UCHL3 may directly regulate the EC-specific function of Aurora B one of the main factors driving fidelity of MT-KT attachments (7, 8, 10–12). First, UCHL3 interacts with Aurora B during prometaphase (Fig. 5C) presumably with the ubiquitin-modified, slower-migrating forms of this kinase. In addition, slower migrating forms of Aurora B accumulate in the UCHL3 knock-out cells specifically during prometaphase (Fig. 5A), when the EC pathway is active. Furthermore, immunoprecipitations under denaturing conditions reveal presence of polyubiquitylated form of Aurora B in the prometaphase cells, a modification which strongly accumulates in the absence of UCHL3 (Fig. 5D). Finally, UCHL3 DUB activity is crucial for the role of UCHL3 in the EC pathway and chromosome bi-orientation (Fig. 3C-E). Together, these results are consistent with the possibility that UCHL3 directly deubiquitylates polyubiquitylated Aurora B during prometaphase.

It is unlikely that UCHL3-mediated deubiquitylation regulates proteolysis of Aurora B as the protein levels of Aurora B remain unchanged through mitotic progression in the absence of UCHL3 (Fig. 5A, B, Fig. S1F). Previous findings demonstrated the role of CUL3-mediated monoubiquitylation of Aurora B (39–42) recognized by the ubiquitin-binding protein UBASH3B (30, 43) in the localization of this kinase to centromeres and microtubules. However, we could not detect any changes in the dynamic localization of Aurora B in UCHL3-downregulated and knock-out cells (Fig. 6A, B, E) and the absence of UCHL3 led to increase in polyubiquitylation and not monoubiquitylation signals on Aurora B (Fig. 5D). Moreover, the levels of NEDD8-modified CUL3 remained unchanged in the absence of UCHL3 (data not shown), excluding the possibility that UCHL3 indirectly affects activity of CUL3, which could be anticipated based on previously reported dual deubiquitylating and deNeddylating activity of UCHL3 (44, 45). While future experiments will have to identify UCHL3-sensitive specific linkage type of polyubiquitin chains detected on Aurora B, it is unlikely that UCHL3 acts in the CUL3-UBASH3B signaling axis and in proteolysis of Aurora B.

Interestingly, our findings suggest that one possible molecular consequence of the increase in Aurora B polyubiquitylation in the absence of UCHL3 is the reduced interaction with the mitotic centromere-associated kinesin (MCAK) but not with other Aurora B-binding partners such as Astrin and the CPC component Survivin (Fig. 6F). MCAK activity as a microtubule depolymerase on incorrectly formed MT-KT attachments is inhibited by direct Aurora B-mediated phosphorylation (14–20) and deletion of MCAK leads to increased number of laterally attached kinetochores resulting in bi-orientation defects and segregation errors (17). We speculate, that UCHL3-driven deubiquitylation of Aurora B promotes its interaction with MCAK allowing for stabilization of MT-KT attachments as cells progressively form metaphase plate (Fig. 6G). This fast and reversible mechanism would allow for fine-tuning of EC machinery during prometaphase progression. While future studies are needed to understand the identity of the E3-ubiquitin ligase able to catalyze this UCHL3-sensitive Aurora B polyubiquitylation, it appears possible that UCHL3-mediated regulation is restricted to the EC correction pathway and does not seem to have an impact on SAC signaling. These results are surprising because knockdown of UCHL3 results in polar chromosomes with only partially attached or completely unattached kinetochores (Fig. 4A-C), a state which should generate a strong SAC response leading to prometaphase arrest. In contrast, our data demonstrate that the recruitment of SAC protein BubR1 to unattached kinetochores occurs normally in UCHL3-deficient cells (Fig. S2A-C). Likewise, the onset of anaphase is not delayed in the absence of UCHL3 (Fig. 2D, H, Fig. S1C, D, F). Since Aurora B plays an essential direct role not only in the EC but also in the SAC pathway for instance through kinetochore recruitment of SAC protein BubR1 (26), our data support a model where UCHL3-mediated deubiquitylation is able to functionally uncouple EC and SAC mechanisms during prometaphase. As such, it offers interesting research perspectives on the EC pathway with unprecedented precision. Since UCHL3 has been recently implicated in human cancer (32, 33), our data also open interesting perspectives for studying the role of EC pathway in the carcinogenesis process.

## Nonstandard Abbreviations

MT-KT: microtubule-kinetochore
SAC: spindle assembly checkpoint
MAD2: mitotic arrest deficient 2
BubR1: Bub1-related 1 kinase
MCC: Mitotic Checkpoint Complex
APC/C: Anaphase Promoting Complex/Cyclosome
EC: error-correction
CPC: Chromosomal Passenger Complex
MCAK: mitotic centromere-associated kinesin
DUBs: deubiquitylases
UCHL3: Ubiquitin carboxyl-terminal hydrolase isozyme L3
UCH: C-terminal hydrolases
BAP1: BRCA1 associated protein 1
TCID: 4,5,6,7-Tetrachloro-1*H*-Indene-1,3(2*H*)- dione
DAPI: 4′,6-Diamidino-2-phenylindole dihydrochloride
PP1*γ*: Protein phosphatase 1 gamma
STLC: S-Trityl-L-cysteine

## Acknowledgments

We thank Snezhana Oliferenko, Manuel Mendoza, Bill Keyes and the members of the Sumara group for helpful discussions on the manuscript. Kallayanee Chawengsaksophak for mentorship and support. We thank the Imaging Center of the IGBMC (ICI) for help on confocal microscopy and to the IGBMC core facilities for their support on this research. K.J. was supported by a fellowship from Gouvernement français et L’Institut français de Prague, Charles University, Prague, a LabEx international PhD fellowship from IGBMC and a fellowship from the “Ligue Nationale Contre le Cancer”. Y.L. is supported by a PhD fellowship from the China Scholarship Council (CSC). Research in R.S. laboratory was supported by funding from The Czech Academy of Sciences (RVO68378050) and from the Czech Centre for Phenogenomics by the Ministry of Education, Youth and Sports of the Czech Republic (LM2015040). This study was supported by the grant ANR-10-LABX-0030-INRT, a French State fund managed by the Agence Nationale de la Recherche under the frame program Investissements d’Avenir ANR-10-IDEX-0002-02. Research in I.S. laboratory was supported by IGBMC, CNRS, Fondation ARC pour la recherche sur le cancer, Institut National du Cancer (INCa), Agence Nationale de la Recherche (ANR), Ligue Nationale contre le Cancer, USIAS and Sanofi iAward Europe.

## Author contributions

K.J. and Y.L. designed and performed experiments and helped writing the manuscript. C.K., S.F., A.A.A. and M.D. helped performing experiments. L.B. helped designing and performing the siRNA screens. R.S. helped designing the experiments and supervising the project. I.S supervised the project, designed experiments and wrote the manuscript with input from all authors. The authors declare no competing financial interests.

## Supplementary Materials

**Fig. S1.**
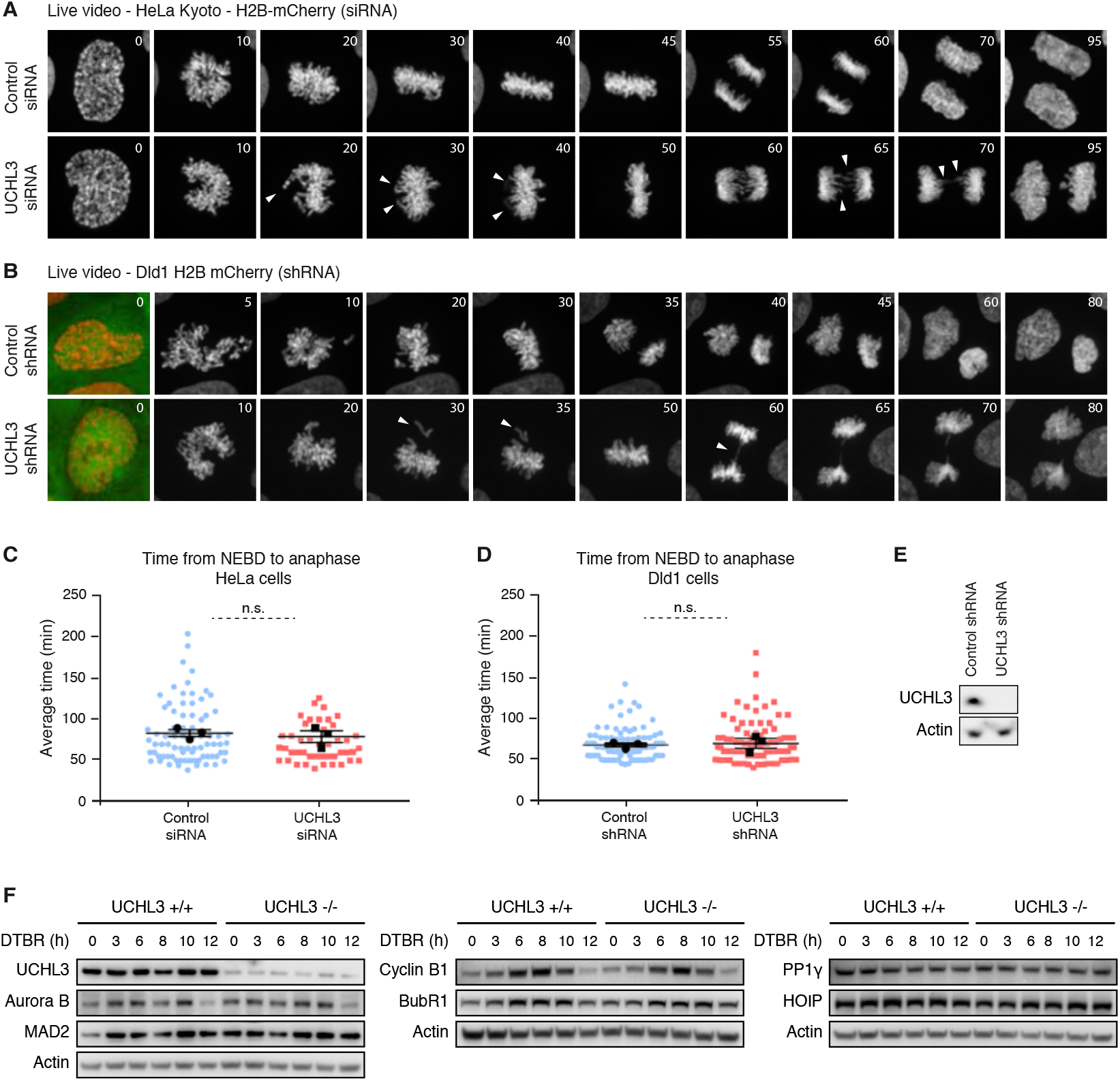
UCHL3 regulates fidelity but not timing of chromosome segregation. (A) HeLa cells stably expressing H2B-mCherry (DNA) and MAD2-LAP were treated with siRNA as indicated and synchronized by double thymidine block and release (DTBR) and analyzed by spinning disk live video microscopy. t=0 corresponds to nuclear envelope breakdown (NEBD). Selected time-frames show DNA channel (gray). Arrowheads point to misaligned chromosomes and lagging chromosomes. (B) Dld1 cells stably expressing H2B-mCherry and GFP-shRNAs as indicated were synchronized by DTBR, t=0 represents NEBD. Selected time frames of Dld1 cells show DNA channel (gray), t=0 is merge of DNA (red) and GFP-shRNA (green). Arrowheads point to misaligned chromosomes and lagging chromosomes. (C) Average time needed from prometaphase to anaphase from cell shown in A, mean (min) ± SEM of 3 experiments. Control = 83.6 ± 4.2, UCHL3 siRNA = 79.4 ± 7.2, p= 0.6433. Colored dots represent values of individual cells, black symbols represent the mean experimental value. (D) Average time needed from prometaphase to anaphase from cells shown in B, mean (min) ± SEM of 3 experiments. Control = 68.0 ± 2.6, UCHL3 shRNA 69.9 ± 6.1, p= 0.7829. Colored dots represent values of individual cells, black symbols represent the mean experimental value. (E) Confirmation of UCHL3 knockdown by western blotting. (F) Western blot analysis showing protein levels of selected proteins upon double thymidine block and release using UCHL3 +/+ and UCHL3 −/− HeLa cells. Numbers indicate the time after release from thymidine in hours.

**Fig. S2.**
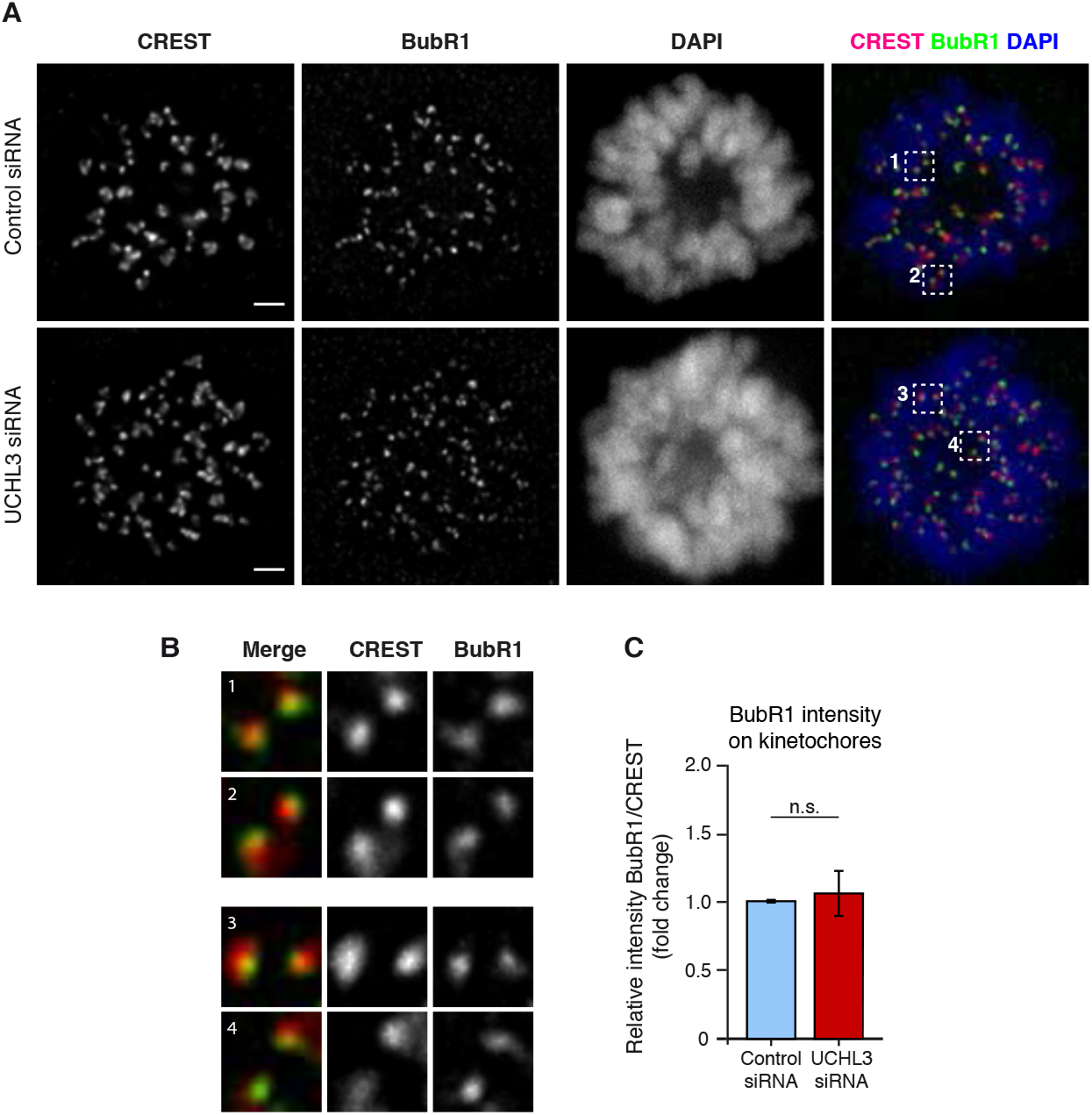
UCHL3 does not regulate SAC signaling. (A) HeLa cells were transfected with indicated siRNAs, synchronized in prometaphase by monastrol and analyzed by immunofluorescence, scale bar 2μm. (B) Magnified regions in numbered boxes are showing co-localization of BubR1 (green) and kinetochore (red) signals. (C) Quantification of relative BubR1 intensity (BubR1/CREST ratio) on kinetochores normalized to the control shown as mean ± SEM of 3 experiments. Control = 1, n= 2740 kinetochores, UCHL3 siRNA 1.06 ± 0.18, n= 2831 kinetochores, p= 0.7765.

**Fig. S3.**
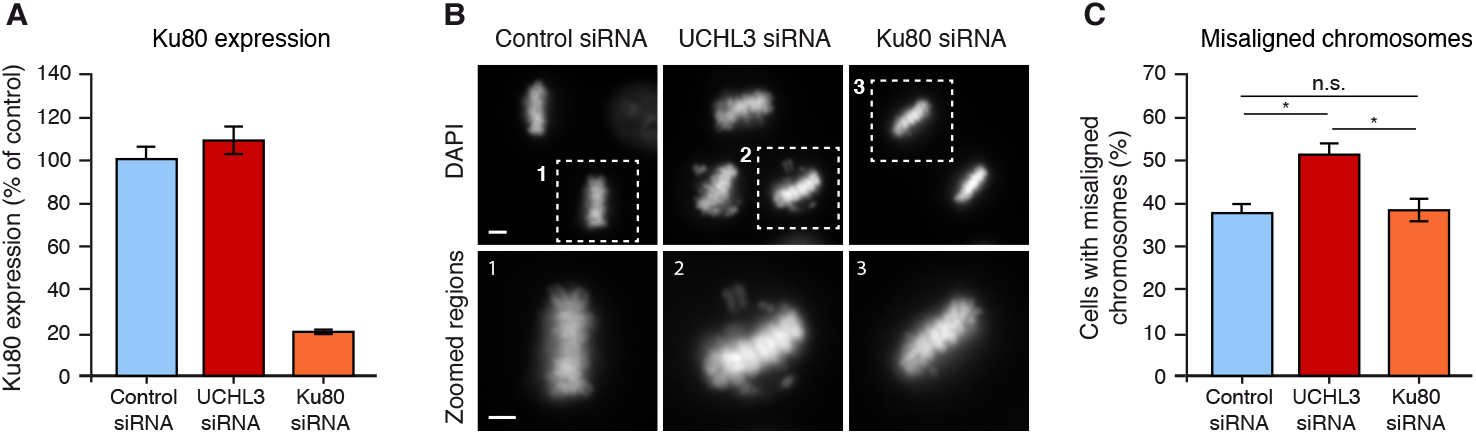
UCHL3-mediated chromosome alignment is Ku80 independent. (A) HeLa cells were treated with indicated siRNAs, synchronized to metaphase and Ku80 expression was analyzed by qPCR. (B) HeLa cells were treated with indicated siRNAs, synchronized to metaphase and analyzed by immunofluorescence. Lower panel shows magnified regions corresponding to numbered boxes, scale bar 2μm. (C) Percentage of cells having at least one misaligned chromosome was counted, mean ± SEM of 3 experiments. Control = 38.2 ± 2.2, n= 318 cells, UCHL3 3’UTR siRNA = 52.1 ± 2.3, n = 326 cells, Ku80 siRNA = 38.4 ± 3.3, n = 350 cells, control to UCHL3 siRNA p = 0.0123, control to Ku80 siRNA p = 0.9570, UCHL3 siRNA to Ku80 siRNA p = 0.0267.

**Table S1.**
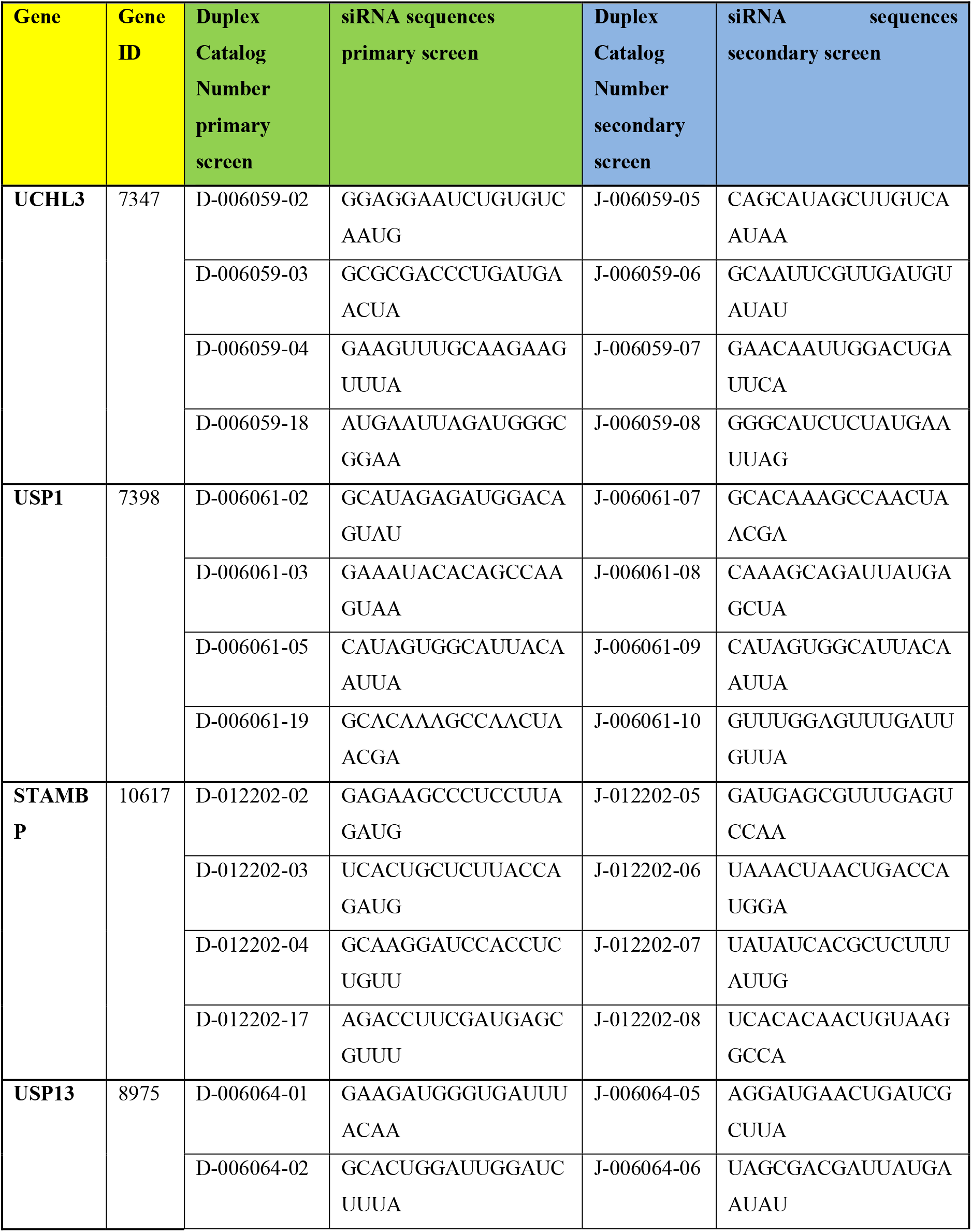
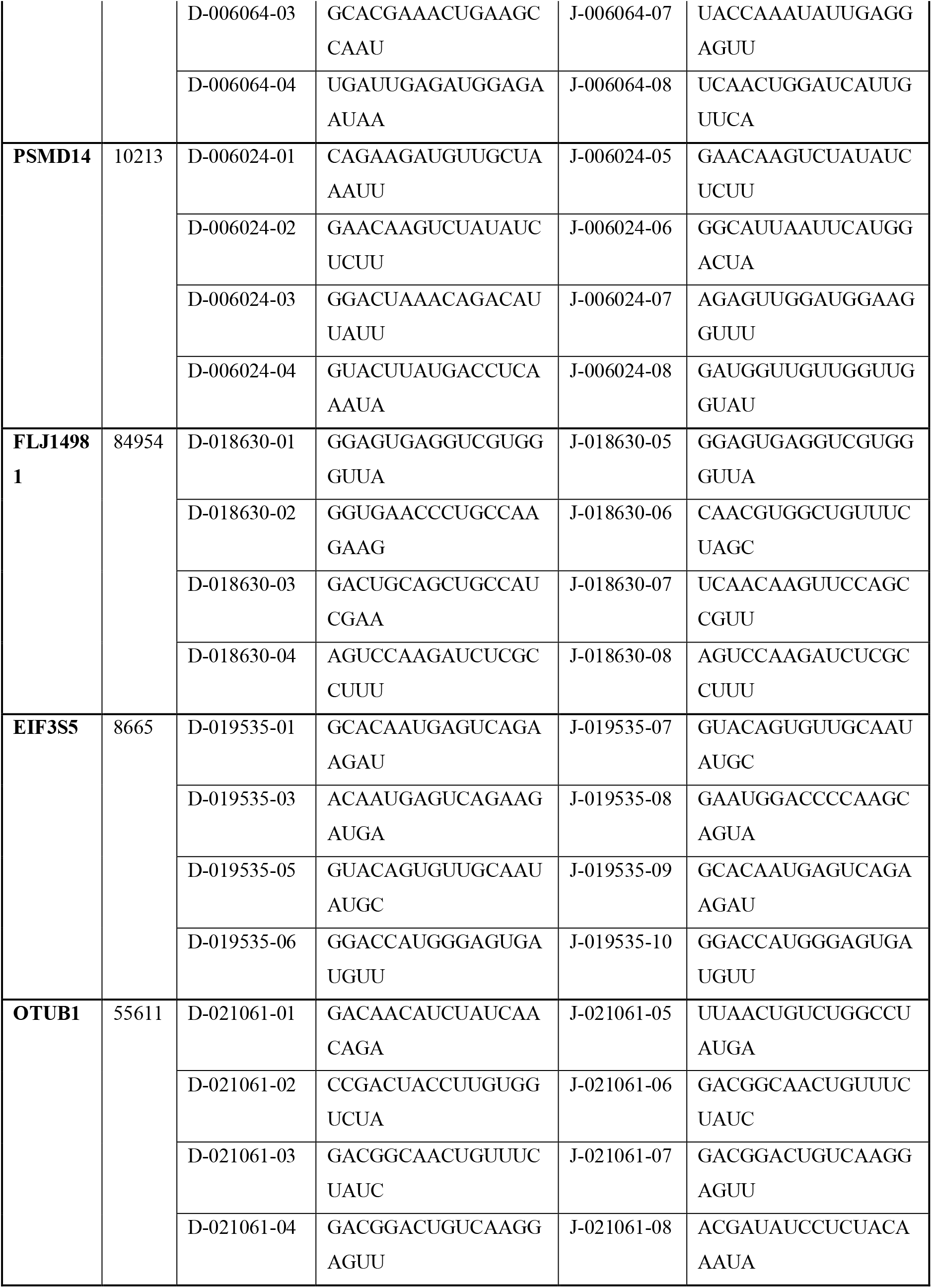
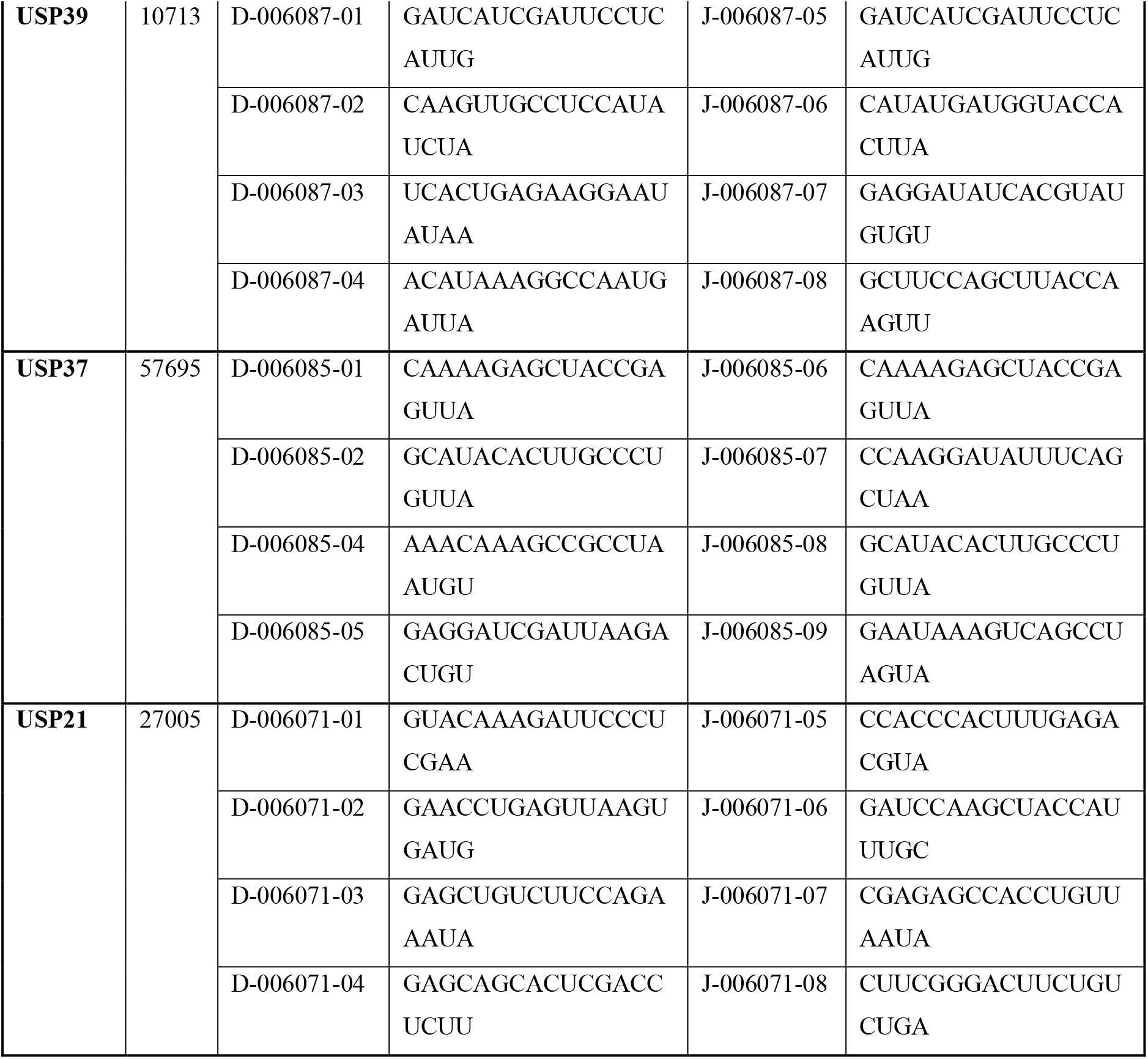
The list of DUB hits in the siRNA screen. The list of DUB genes identified by the primary and secondary siRNA screens. For each hit the gene symbol, gene ID are indicated. The list includes the individual siRNA oligonucteotides of the siGENOME library used as pools (SMARTpools) for targeting indicated genes (Gene Symbol and Gene ID) in the primary siRNA screen. The Dharmacon pool catalog numbers, duplex catalog numbers and the sequences are indicated. For the secondary siRNA screen, the list of the individual siRNA oligonucteotides of the ON-TARGETplus library used as pools (SMARTpools) for targeting the indicated genes (Gene Symbol and Gene ID) The Dharmacon pool catalog numbers, duplex catalog numbers and the sequences are indicated.

